# Fine-scale characterization of genomic structural variation in the human genome reveals adaptive and biomedically relevant hotspots

**DOI:** 10.1101/294322

**Authors:** Yen-Lung Lin, Omer Gokcumen

## Abstract

Genomic structural variants (SVs) are distributed nonrandomly across the human genome. These “hotspots” have been implicated in critical evolutionary innovations, as well as serious medical conditions. However, the evolutionary and biomedical features of these hotspots remain incompletely understood. In this study, we analyzed data from 2,504 genomes from the 1000 Genomes Project Consortium and constructed a refined map of 1,148 SV hotspots in human genomes. By studying the genomic architecture of these hotspots, we found that both nonallelic homologous recombination and non-homologous mechanisms act as mechanistic drivers of SV formation. We found that the majority of SV hotspots are within gene-poor regions and evolve under relaxed negative selection or neutrality. However, we found that a small subset of SV hotspots harbor genes that are enriched for anthropologically crucial functions, including blood oxygen transport, olfaction, synapse assembly, and antigen binding. We provide evidence that balancing selection may have maintained these SV hotspots, which include two independent hotspots on different chromosomes affecting alpha and beta hemoglobin gene clusters. Biomedically, we found that the SV hotspots coincide with breakpoints of clinically relevant, large *de novo* SVs, significantly more often than genome-wide expectations. As an example, we showed that the breakpoints of multiple large *de novo* SVs, which lead to idiopathic short stature, coincide with SV hotspots. As such, the mutational instability in SV hotpots likely enables chromosomal breaks that lead to pathogenic structural variation formations. Our study contributes to a better understanding of the mutational landscape of the genome and implicates both mechanistic and adaptive forces in the formation and maintenance of SV hotspots.

## Introduction

Structural variants (SVs) are an important source of polymorphic genomic variation within primate species (Gokcumen et al. 2011, 2013a; Sudmant et al. 2015; Conrad et al. 2009). Unlike single nucleotide variants, SVs involve blocks of sequences that vary in copy number (copy number variants) (Conrad et al. 2009), chromosomal locations (translocations), or directionality (inversions) (Feuk et al. 2006; Alkan et al. 2011). Given their larger size, an individual SV potentially has a higher phenotypic impact than single nucleotide variants, especially when affecting functional sequences (Conrad et al. 2009; Iskow et al. 2012a, 2012b). In fact, SVs have been associated with many human diseases (Weischenfeldt et al. 2013), especially autoimmune, metabolic and cognitive disorders (Polley et al. 2015; Hollox et al. 2008; Stefansson et al. 2008; Traherne et al. 2010). However, the haplotype architectures around SVs are generally complex due to repeat content, gene conversion and recurrent formation of SVs, making it difficult to accurately impute them in genome-wide association studies (Sudmant et al. 2015). As such, it is plausible that many disease-associated SVs are yet to be revealed. For example, recent locus-specific studies have unearthed novel SVs with associations to schizophrenia (Sekar et al. 2016) and blood cholesterol levels (Boettger et al. 2016) that were invisible to previous genome wide association studies. So, the biomedical impact of SVs is likely greater than is currently appreciated.

SVs exhibit a nonrandom distribution across the human genome. For example, it has been observed that SVs cluster into “hotspots” on primate genomes (Perry et al. 2006; Gokcumen et al. 2011). These early studies have developed both mechanistic and evolutionary scenarios to explain the formation and maintenance of these hotspots. Specifically, Perry et al. (2006) highlighted non-allelic homologous recombination during meiosis as the major mechanistic contributor to hotspot formation. Gokcumen et al. (2011) identified non-neutral evolutionary forces that have maintained hotspots since before the human-rhesus macaque ancestor. However, the recent and more complete human SV datasets (Sudmant et al. 2015) allow us to answer three questions in a more definitive manner: What are the features of local genomic architecture facilitating the formation of hotspots? What are, if any, the adaptive forces that maintain hotspots? What is the biomedical impact of SV hotspots in human genome?

The presence of SV hotspots can be understood as an interplay between the mutational mechanisms that give rise to them and the evolutionary forces that eliminate or maintain them. Indeed, previous studies have reported the coincidence of SVs with specific genomic architectural features (Hastings et al. 2009). One of the genomic features commonly accompanying SVs is segmental duplication content (Sharp et al. 2005; Bailey and Eichler 7/2006; She et al. 2008). The long and highly homologous sequences of tandem segmental duplications enable the misalignment between paralogous counterparts during recombination, which leads to unequal crossing over. This process is called non-allelic homologous recombination, or NAHR. Segmental duplication-mediated NAHR is a major cause of recurrent, *de novo* genomic rearrangements that are associated with rare disorders (Dittwald et al. 2013). Non-pathogenic SVs are also commonly observed within segmental duplication rich regions (Conrad et al. 2009). Segmental duplications are discussed as the primary genomic element that facilitates the formation of SV hotspots (Hastings et al. 2009), especially in great apes, the genomes of which have been shaped by a “burst” of segmental duplications (Gokcumen et al. 2013a; Marques-Bonet et al. 2009).

Another component of genomic architecture associated with SVs is transposable elements. For example, artificially introducing a single copy of the highly repeated Ty element into the yeast genome increased the local rate of genomic rearrangements (Chan and Kolodner 2011). In primate genomes, the rearrangements mediated by retroelements were suggested as an important factor in primate genome evolution. Moreover, studies have discovered cases of human SVs generated through Alu-, HERV-, or LINE-mediated NAHR (Robberecht et al. 2013; Campbell et al. 2014; Startek et al. 2015). However, the relationship between transposable elements and SV hotspots has not been directly interrogated.

Mechanisms other than recombination-based errors (*e.g.*, NAHR) and transposable element activity are also significant contributors to structural variation formation (Hastings et al. 2009). These include, but are not limited to non-homologous events, such as non-replicative non-homologous repair, replication slippage, fork stalling and template-switching. These events are harder to map and often can lead to complex rearrangements involving multiple SVs (Zhao et al. 2016). To further complicate the issue, the individual mechanisms are not necessarily independent of each other and the genomic architectural features that facilitate some of these mechanisms are not uniformly distributed across the genome. For example, Alu-richness in a locus was argued to facilitate structural variation formation through both recombination- and repair-based mechanisms (Boone et al. 2014). The exact impact of these mechanisms across the genome and their codependency is not precisely known.

While the formation of SVs depends on their local architectures, their maintenance depends on the evolutionary forces acting upon them. It is suggested that SVs that disrupt functional sequences are mostly deleterious, and therefore have been eliminated from the human population via negative selection (Conrad et al. 2005) or drift. Indeed, previous studies have shown a drastic depletion of deletion polymorphisms from exonic regions (Conrad et al. 2009). Despite the overall depletion of SVs in functional sequences, there are still some SVs that are passively tolerated under relaxed negative selection (Nguyen et al. 2008; Eaaswarkhanth et al. 2016), and some others that are actively maintained by non-neutral evolutionary forces (Gokcumen et al. 2013b; Pajic et al. 2016; Perry et al. 2007; Polley et al. 2015).

In this paper, we use recently available, high-resolution datasets (Sudmant et al. 2015) to define and investigate SV hotspots in the human genome both from mechanistic and evolutionary perspectives, and investigate their biomedical impact.

## Results

### High-resolution detection of hotspots confirms non-random distribution of SVs in the human genome

We determined the SV hotspots by dividing the human genome (hg19) into fixed-size, non-overlapping intervals, and directly counting the number of SVs in each interval. Using intervals of fixed size simplified comparisons of genomic content (e.g., gene content, GC content, etc.) among intervals.

After conducting preliminary analyses involving multiple interval sizes, we decided to use 100 kb intervals (28,103 intervals across the genome, **Table S1**). This served to avoid the majority of situations where larger SVs spanned multiple intervals. To reduce bias when determining the SV hotspots, our dataset excluded intervals that overlapped with sequencing gaps. Such gaps can contribute to the underestimation of SV number.

Overall, we worked with 42,758 SVs (deletions, duplications, multiallelic copy number variants, inversions, and insertions) called from 2,504 human genomes across 26 populations available through the 1,000 Genomes Project Phase 3 data release (1000 Genomes Project Consortium et al. 2015). To our knowledge, it is the most accurate SV map for healthy humans (Sudmant et al. 2015). Moreover, the same data release also provides the best annotated single nucleotide variation for thousands of human genomes. Therefore, it provides an unprecedented inclusiveness, accuracy, and genomic context to the study of hotspots of structural variation. It is important to note that this dataset aims to be very accurate (Sudmant et al. 2015), but sacrifices from sensitivity for achieving this accuracy.

Prior studies have demonstrated that SV distribution across the genome is nonrandom (Perry et al. 2006; Gokcumen et al. 2011). To replicate this finding in our interval-based framework, we first built a Poisson probability distribution, where we assumed that the 42,758 SVs fall randomly into the 28,103 intervals (**Figure 1**). This allowed us to model the range of expected SV count within an interval. We then compared the empirical observation (mean=1.83, sd=1.84) to the expected distribution (mean=1.84, sd=1.35). We found that there are 1,148 intervals (4% of all intervals) that harbor 6 or more SVs, which put them at the 99^th^ percentile or higher estimated from the expected distribution of SV numbers in each interval (hotspots, **Table S2**). This is significantly higher than expected by chance under the assumption of a random distribution (**p<0.0001, Chi-square test**). Conversely, we also found that the number of intervals where we found no SVs (i.e., deserts, **Table S2**) is also higher in the observed data (6,827) as compared to the expected distribution (**p<0.0001, Chi-square test**). These results further support the nonrandom distribution of SVs across the human genome (**Figure 1**).

**Figure 1.**
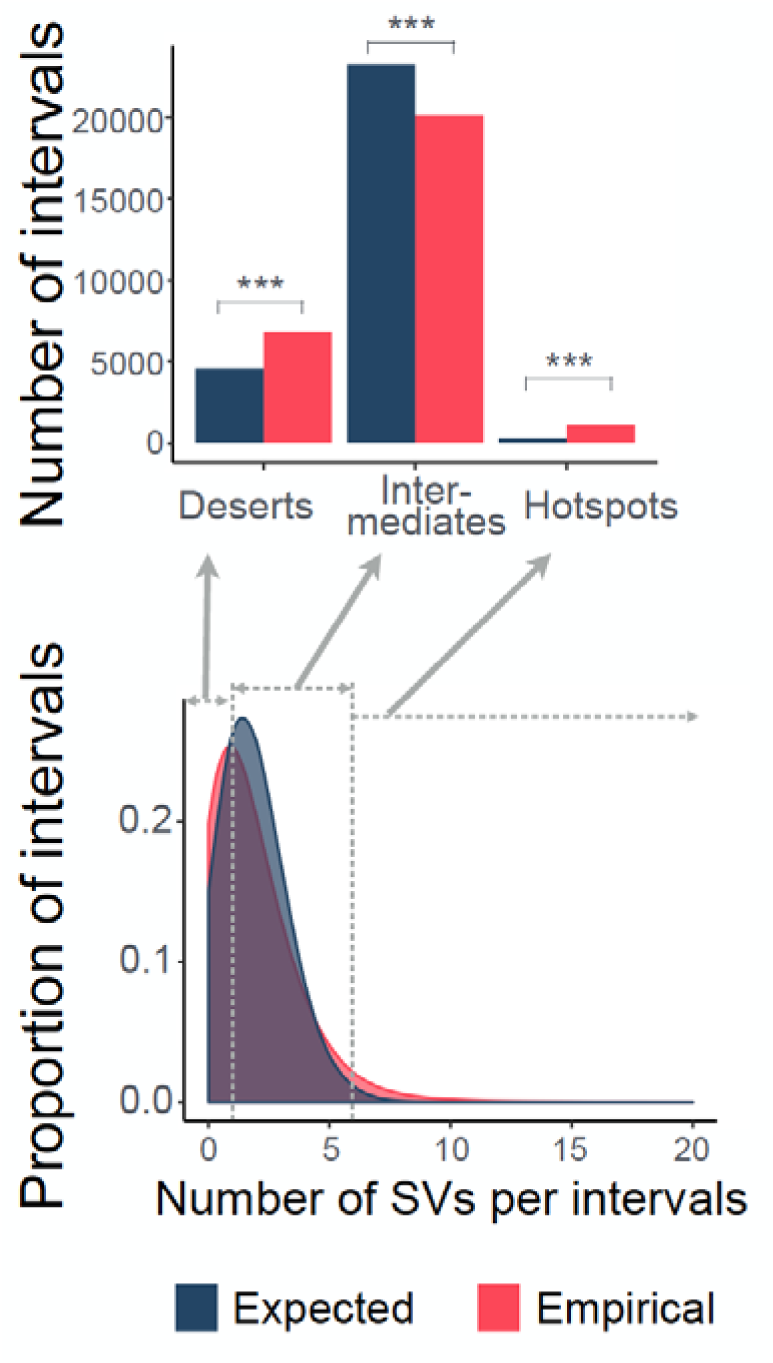
The nonrandom distribution of SVs across human genomes. The density plot (lower panel) overlay the empirical and the expected probability distribution of the number of SV(s) within an 100Kb-long interval of human reference genome (hg19). The empirical probability distribution was estimated by the actual number of SVs within each of the 28,103 intervals. The expected probability distribution was estimated assuming a Poisson distribution. As compared to the expected data, the empirical data have higher proportions for both SV deserts (with no SVs) and SV hotspots (with 6 or more SVs). The upper panel shows the significant enrichment of SV deserts and SV hotspots in the empirical data. *** indicates where there is a significant difference with P-values smaller than 0.05 as calculated by Chi-square test.

### Non-allelic homologous recombination and non-homologous mechanisms, but not mobile element content, coincide with SV hotspots

Previous studies have shown that SV hotspots coincide with segmental duplications (Perry et al. 2006). Non-allelic homologous recombination (NAHR) facilitated by segmental duplications leads to recurrent genomic rearrangements (Sharp et al. 2005), which in turn contribute to the formation of SV hotspots (Perry et al. 2006). To replicate this finding, we tested the hypothesis that SV hotspots are enriched for segmental duplications. To avoid biases in this correlation, we randomly matched non-hotspots intervals with specific hotspot intervals for their genomic composition (i.e., GC content, number of base pairs attributed to coding sequence, mobile elements, and single nucleotide variation, **Table S3**). Using this matched dataset, we were able to replicate the observation that the segmental duplication coverage of the hotspot intervals is significantly higher than the segmental duplication coverage of the matched non-hotspot intervals (**Figure 2A**, **p=9.962×10^−4^, Mann-Whitney U test**). This observation implicates that NAHR mediated by segmental duplications, which present in ~30% of the SV hotspots, is a major mechanism that lead to the formation of SV hotspots. In other words, segmental duplications increase the SV mutation rate in these hotspot regions (Liu et al. 2011). However, it is important to note here that there are still ~70% of the SV hotspots whose formation cannot be explained by segmental duplication-mediated NAHR.

**Figure 2.**
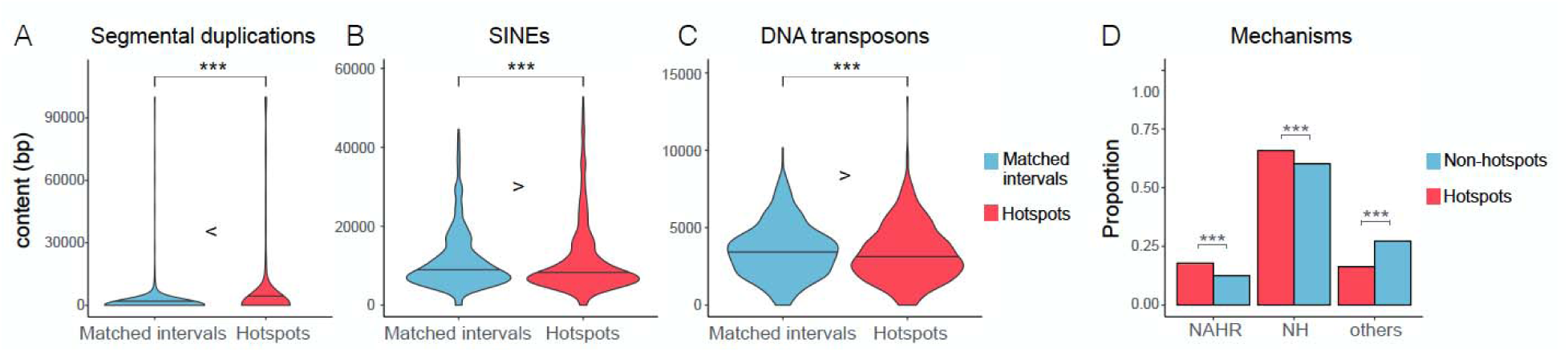
The architectural characteristics and SV formation mechanisms of SV hotspots. The violin plots compare the number of base pair(s) constituting segmental duplications (A), SINEs (B), and DNA transposons between the hotspots (red) and matched non-hotspot intervals (blue). The horizontal line on the plots indicate the median values. The *** signs indicate where there is a significant difference with a P-value lower than 0.05, calculated by Mann-Whitney U test. Panel (D) illustrates the coincidence of NAHR, NH, and other SV formation mechanisms of the SVs within (red) or outside (blue) of hotspots. For each category, the proportions of SVs attributed to individual mechanisms are plotted. *** indicates where there is a significant difference with a P-value smaller than 0.05, calculated by Fisher’s exact test.

We then asked whether mechanisms other than segmental duplication-mediated NAHR are correlated with SV hotspots. One obvious candidate is mobile elements. Mobile elements can generate local genomic instability in many ways. For example, the homologous sequences between the dispersed mobile elements can serve as substrates for NAHR and lead to large genomic rearrangements (Cordaux and Batzer 2009; Kazazian and Goodier 2002). To test the involvement of fixed mobile elements in SV hotspot formation, we compared the SV hotspots and their matched, non-hotspot intervals (**Table S3**) for their composition of fixed LINEs, SINEs, and DNA transposons (as reported in RepeatMasker (Smit et al. 2016)). To our surprise, we did not find any enrichment of mobile elements in the hotspots. In fact, we observed a slight but significant depletion of SINE elements and DNA transposons in SV hotpots as compared to the matched non-hotspot intervals (**Figure 2B**, **SINE: p=0.04419; Figure 2C, DNA transposon: p=0.01719, Mann-Whitney U test**).

This finding contradicts the prediction from individual cases of SVs driven by retroelement-mediated NAHR (Belancio et al. 2009) and other observations where Alu richness leads to recurrent SV formation (Boone et al. 2014). We were especially surprised that LINE elements are not significantly enriched in human SV hotspots despite their large size (**Figure S1A**). To further interrogate this finding, we used available Drosophila data (Zichner et al. 2013) to conduct a similar analysis. We found a significant enrichment of Drosophila mobile element content in the Drosophila hotspots (defined the same way) as compared to the non-hotspot intervals (**Figure S1B**, **p<2.2×10^−16^, Mann-Whitney U test**), contradicting our observations in the human genome. This disparity can be explained, at least partially, by the observation that Drosophila LINE elements are overall much larger and less degraded than those observed in humans. Regardless, our results showed that even though retrotransposons can induce SV formation in principle, rather short and differentiated fixed human retrotransposons are not a major contributor to SV hotspot formation in humans.

Non-homologous (NH) mechanisms potentially contribute to formation of SV hotspots. These include non-homologous end-joining (NHEJ) events and template-switching during replication (Hastings et al. 2009). This type of mechanism does not show dependency on recombination and is difficult to identify based on the context of local architecture alone (Lam et al. 2010). As such, we used a previous study that has categorized a subset of SVs across the genome based on their formation mechanisms (Abyzov et al. 2015). Using this dataset, we found that NH mechanisms, along with NAHR, contributed significantly to SV hotspots (**Figure 2D, p=0.00164, Fisher’s exact test**). Collectively, our results indicate that rather than overactivity of a particular mechanism, both NAHR and NH-based mechanisms contribute to SV hotspots. Therefore, mechanistic disposition of a region for *de novo* SV formation (e.g., presence or absence of segmental duplications) does not fully explain the presence and distribution of SV hotspots in the human genome.

### Negative selection shapes the distribution of hotspots in the genome

Previous studies have suggested that negative selection against SVs is the predominant selective force that shapes the distribution of SVs in the genome (Conrad et al. 2009). As such, we hypothesize that SV hotspots would be biased away from conserved regions. Indeed, when we compared our SV hotspots to regions that are matched for other genetic features (**Table S3**), we found that hotspots are significantly depleted for both coding and genic sequences (**coding sequence: Figure 3A, p=1.673×10^−7^; genic sequences: Figure S2, p=0.002901, Mann-Whitney U test**). In addition, we found a significant depletion of DNase I hypersensitive sites, which are enriched for regulatory sequences, in hotspots as compared to the rest of the genome (**Figure S4, p=0.048, Mann-Whitney U test**). These observations are especially relevant within the context of the recent study by Fudenberg and Pollard (Fudenberg and Pollard 2018) suggesting that chromatin properties and their regulatory involvement contribute to evolutionary constraints on the distribution of SVs across the genome.

**Figure 3.**
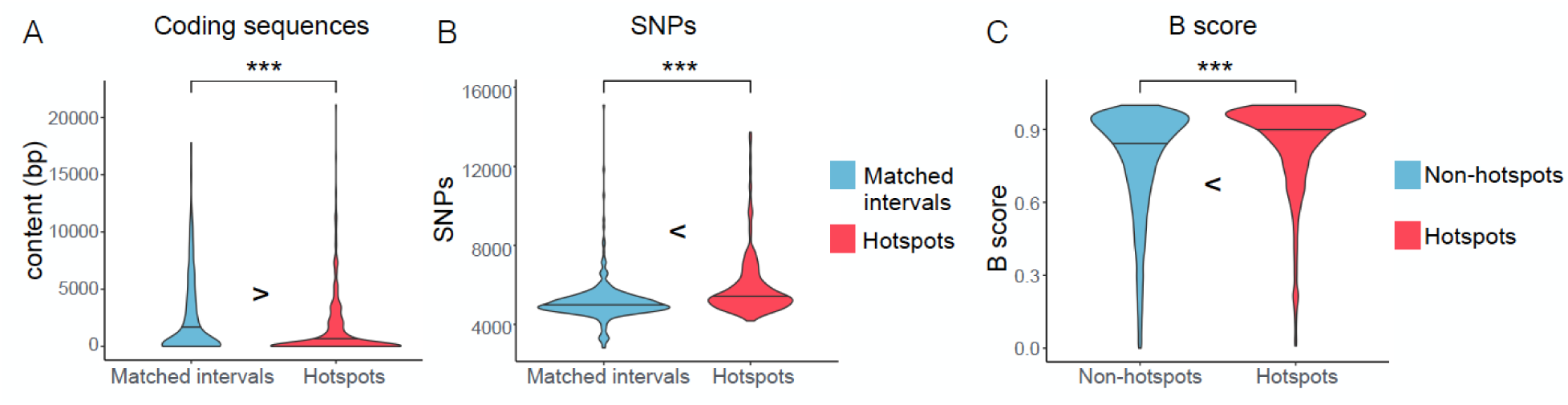
Signatures of relaxed purifying selection in hotspots. Comparisons between hotspots (red) and matched non-hotspot intervals (blue) reveal a significant depletion of the coding sequence content (A), and a significant enrichment of SNPs (B) in the SV hotspots. Panel (C) shows that the hotspots (red) have significantly higher B-statistics as compared to the non-hotspot intervals (blue). B-statistics is a measure of background selection where the higher the B-statistics, the lower the background selection. The horizontal lines indicate the median values. The *** signs indicate where there is a significant difference with a P-value lower than 0.05, calculated by Mann-Whitney U test.

If SV hotspots affect neutrally evolving regions, we also expect an enrichment of single nucleotide variants in the same regions. Indeed, we found that the SV hotspots harbor more single nucleotide variants as compared to the matched non-hotspot intervals (**Figure 3B**, **p < 2.2×10^−16^, Mann-Whitney U test**). With the same logic, we reasoned that even though we found a depletion of fixed SINE elements and DNA transposons as reported above, we still expect an enrichment of polymorphic mobile elements in the hotspot regions. Our results confirmed this expectation as we showed that the SV hotspots are enriched with the insertions of Alu and L1 as compared with the non-hotspot regions (**Figure S2, Alu: p=0.01916; L1: p=0.01668, Mann-Whitney U test**).

These observations all support the hypothesis that negative selection is the primary force shaping the distribution of SV hotspots in the genome. In other words, SV hotspots that overlap with functional sequences have been eliminated by negative selection, while the hotspots overlapping with evolutionarily less conserved sequences remain. To further test this scenario, we used B-statistics, a measure of background selection (McVicker et al. 2009), to demonstrate that SV hotspots overlap with sequences affected significantly less by background selection as compared to the rest of the genome (**Figure 3C**, **p < 2.2 × 10^−16^, Mann-Whitney U test**). SV hotspots, with some exceptions that we discuss below, are distributed away from conserved, phenotypically relevant sequences, supporting the notion that most SV hotspots coincide with neutrally evolving sequences with little fitness effect. This result also suggests that the mutation rate for SVs across the human genome may be relatively high, i.e., high enough to generate hotspots unless eliminated by negative selection.

### Balancing selection maintains the few functionally relevant SV hotspots involving in environment interaction

Despite the general depletion of coding sequences within SV hotspots, some of the SV hotspots harbor a relatively high proportion of genic sequences (**Figure S2A**). To investigate the functional inference of SV hotspots, we conducted functional enrichment analysis on the genes as well as on the putative regulatory sequences harbored in SV hotspots. First, we extracted the protein coding sequences that fall into SV hotspots and conducted a gene ontology enrichment analyses using GOrilla (Eden et al. 2009). We found that blood oxygen transport, sensory perception to smell, synapse assembly, and antigen binding categories are overrepresented among the genes in SV hotspots (**Figure 4A**, **Table S4, Figure S3)**. Some of the specific genes involved in these categories have been reported to be under rapid evolution either because of relaxation of evolutionary constraint and/or positive selection (Voight et al. 2006; Hasin-Brumshtein et al. 2009; Lek et al. 2016). In addition, gene families in these hotspots happen to be also discussed within the context of adaptive evolution (López de Castro et al. 1982; Erlich et al. 1986; Hamza et al. 2010; Krause and Pestka 12/2015; Hollox et al. 2008; Traherne et al. 2010).

**Figure 4.**
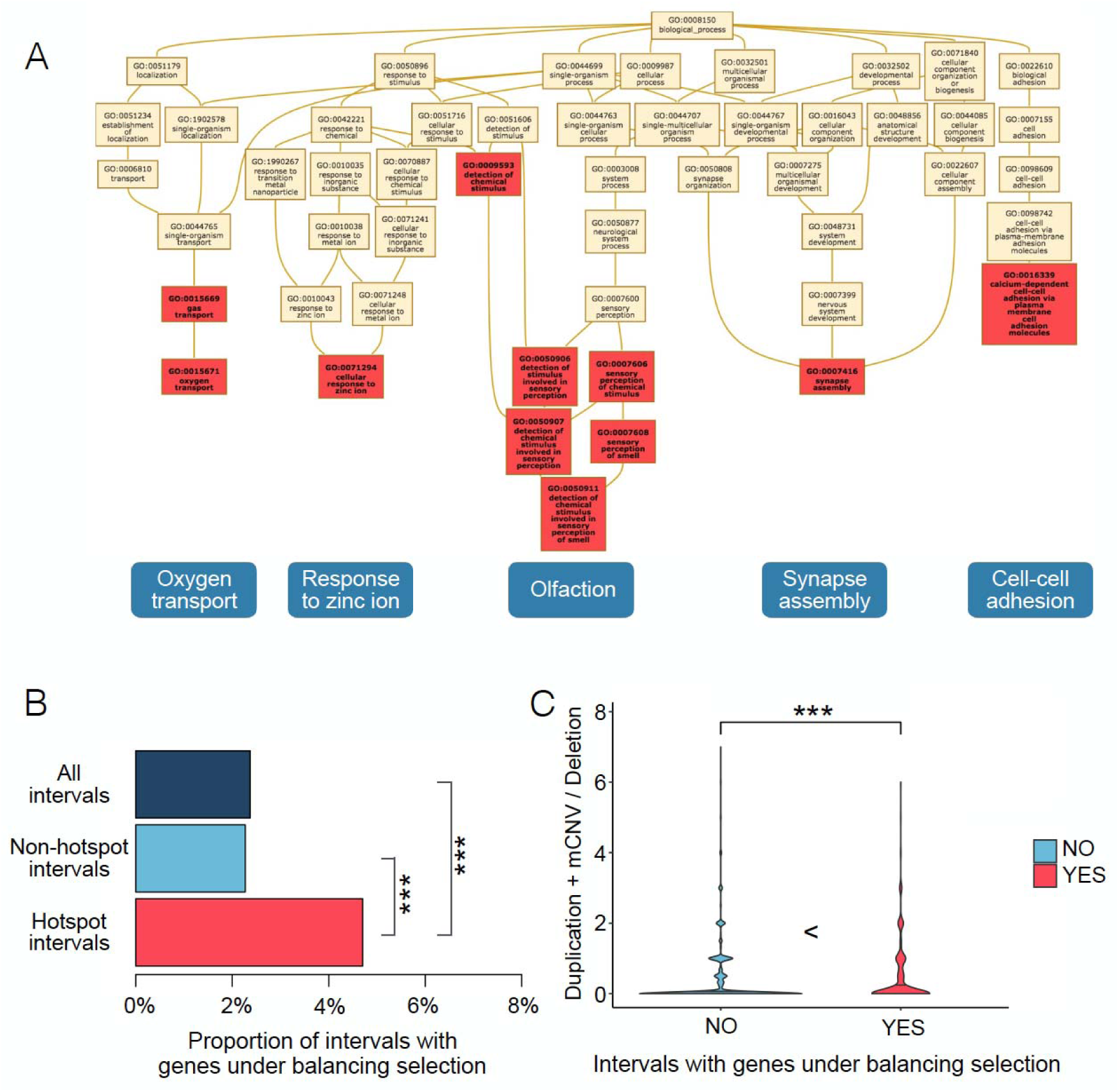
The non-neutral forces and the protein-coding roles of SV hotspots. (A) demonstrates the biological process ontologies overrepresented among the genes with their coding sequences overlapping with hotspots. The overrepresented terms (FDR q-value < 0.05) are highlighted in red. These terms falls into the five distinct categories boxed at the bottom of the figure. (B) shows the proportions of all intervals, the non-hotspot intervals, and the hotspot intervals that overlaps with genes reported as under balancing selection. A significantly higher proportion of hotspot intervals harbor genes under balancing selection, compared to other non-hotspot intervals, as well as the genomic background. The *** signs indicate where there is a significant difference with a P-value lower than 0.05, calculated by Fisher’s exact test. (C) compares between intervals with (red) and without (blue) genes under balancing selection for their ratios of duplication and mCNV variant(s) over deletion variant(s) within an interval. The *** signs indicate where there is a significant difference with a P-value lower than 0.05, calculated by Mann-Whitney U test.

To retrieve a comprehensive picture of the functional impact of SV hotspots, we further look into the potential *cis* functions of the non-coding sequences overlapping with SV hotspots. We used the genomic regions enrichment of annotations tool (GREAT), which associates the non-coding sequences to the genes they potentially regulate (McLean et al. 2010). This tool extends a gene’s annotation from the coding sequence itself to its nearby sequences, and performs over-representation analysis of multiple ontologies (**Table S5**). Complementing the GO analyses described above, we found from the GREAT analysis an overrepresentation of sequences that are involved in olfactory signaling and acquired immunity. In addition, we found that some sperm flagellum-related functions are overrepresented for the sequences harbored in the SV hotspots, concordant with the previous finding that the enhancers active in the male reproductive system are among the enhancers with the lowest evolutionary constraint (Huang et al. 2017). Furthermore, we found that sequences related to pigmentation, both via melanin biosynthesis and melanosome organization, are enriched in SV hotspots regions, thus linking the SV hotspots with partial albinism, iris hypopigmentation, red hair, and cutaneous photosensitivity (**Table S5**).

To explain the presence of such genic hotspots, we propose two mutually inclusive scenarios: First, it is plausible that these SVs are mostly duplications, which leads to gain-of-function rather than loss-of-function. As such, their evolutionary impact may be less likely to be negative as compared to deletion variants. The second possibility is that there is adaptive context, where multiple alleles of the affected sequences are beneficial and actively maintained.

To test these scenarios, we examined the overlap between SV hotspots and a list of 195 genes that show signatures of balancing selection as described in a previous paper (DeGiorgio et al. 2014). We found that genes that were found to be evolving under balancing selection in this dataset are more often located within SV hotspots than expected by chance (**Figure 4B, p=2.176×10^−6^, Fisher exact test**). In parallel, we analyzed the correlation of the proportion of duplications and multiallelic copy number variants as compared to deletions in hotspots. We found that the SV hotspots with genes evolving under balancing selection are also significantly enriched for duplications and multi-allelic copy number variants (**Figure 4C, p=0.006416, Fisher exact test**). These results suggest that balancing selection is a major factor in maintaining SVs in a small subset of SV formation hotspots, especially involving multiallelic copy number variants and duplications,

Combined, our results reveal that the functional sequences within SV hotspots are involved in environmental interaction -blood groups, immunity, perception, and skin function. These findings are consistent with our proposed evolutionary scenario where the relatively small number of functionally relevant SV hotspots are primarily maintained through balancing, diversifying or geography-specific positive selection (Key et al. 2014). These regions should be of great interest in understanding complex diseases, particularly those related to immune-related disorders (**Table S1**).

#### Of hotspots, malaria and blood disorders

To exemplify our findings with regards to balancing selection in SV hotspots, we focused on the alpha and beta globin clusters, both of which are SV hotspots despite being on different chromosomes (**Figure 5**). We are particularly interested in these regions because the variation herein has long been associated with protection against malaria in a balancing selection context (Hill et al. 1991; Modiano et al. 2001). When we investigated the variation in these loci, we found that there are seven distinct SVs in each of the two loci. Almost all known alpha and beta globin genes are influenced by at least one SV. Notably, it is plausible that there are even more SVs in this repeat-rich region which are not present in the 1,000 Genomes dataset, as the project aims to maximize accuracy over sensitivity (Sudmant et al. 2015).

**Figure 5.**
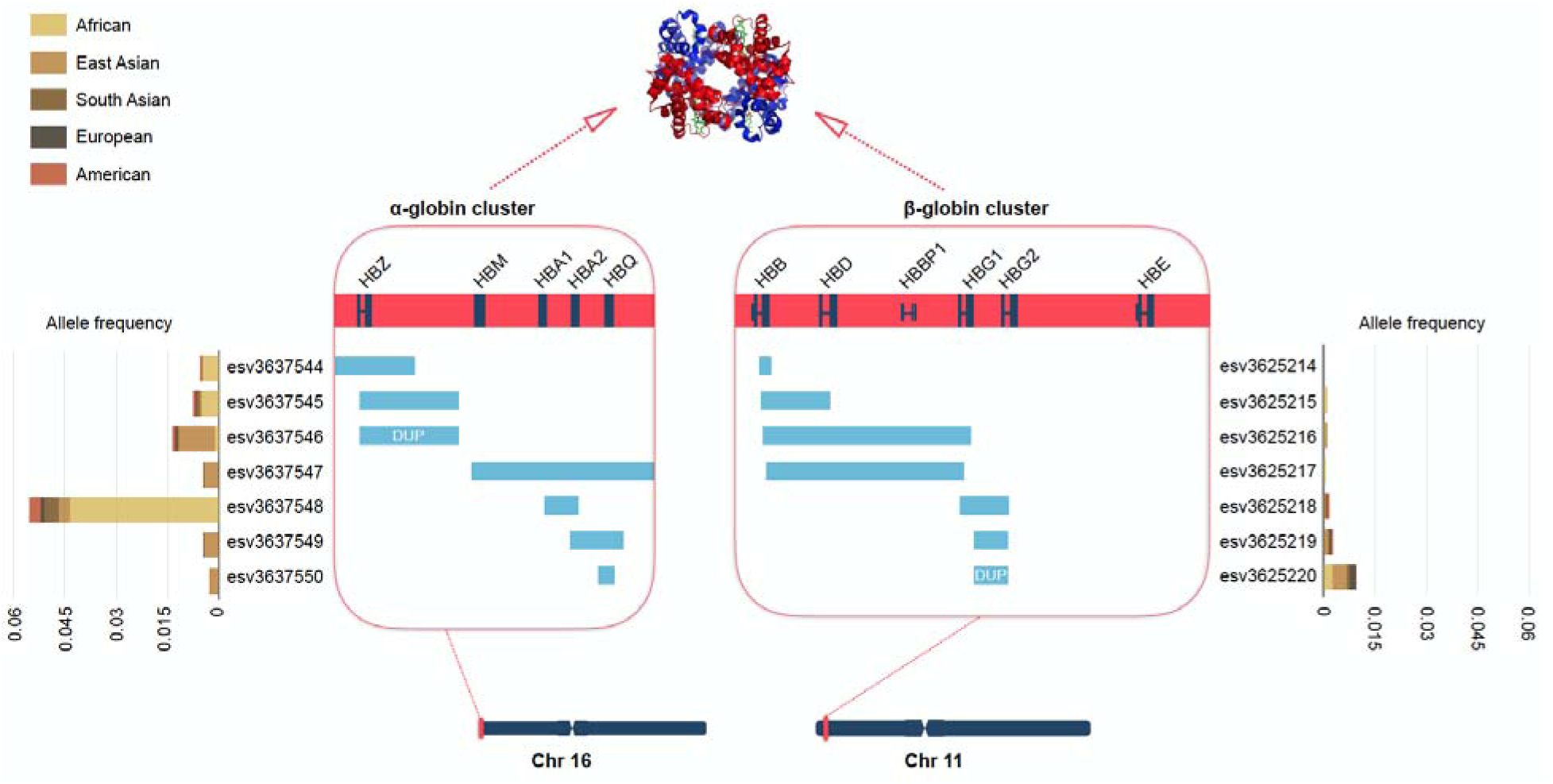
Alpha and beta globin gene clusters are both overlapping within SV hotspots. The figure shows the alpha globin cluster and beta globin cluster on chr16 and chr11. Genes in these two clusters code for the different parts of the hemoglobin protein indicated above -two alpha (red) and the two beta (blue) globins of the hemoglobin tetramer. Within the boxes are enlarged views of the two gene clusters. Genes are plotted in navy across the red bar. Deletion variants are represented as blue bars and duplication variants are blue bars marked with “DUP”. The allele frequency and population composition of individual SVs are shown with the stacked bars next to the SVs.

Some of the SVs in these regions are common even though they overlap with coding exons. These include esv337548, which reaches an allele frequency of 16% in sub-Saharan African populations, where malaria is geographically prevalent. Moreover, this particular deletion has been associated with thalassemia (Embury et al. 1980), which, as a blood disorder with similarities to sickle cell anemia, has been associated with protection against malaria (Clegg and Weatherall 1999; Allen et al. 1997). In other words, esv337548 is a prime candidate to be evolving under balancing selection in a relatively straightforward framework. Briefly, a person carrying the esv337548 deletion homozygously suffers from thalassemia traits, but gains protection against malaria.

Given that the heterozygous genotype does not cause thalassemia and still provides protection against malaria, it is the fittest genotype in the region. This heterozygote advantage has likely maintained both the deleted and non-deleted alleles in the population. The added complexity arises from the fact that other, recurrent SVs in the region have been reported, some of which are also associated with thalassemia (Harteveld et al. 2005, 2008). In fact, the link between malaria, blood disorders, and SV hotspots can further be generalized as other regions that are associated with protection against malaria were found to be SV hotspots in our study, including the *GYPB* locus (Leffler et al. 2017).

### SV hotspots coincide with breakpoints of clinically relevant *de novo* SVs

We further investigated co-occurrence of biomedically relevant variations with hotspots. We compared the number of single nucleotide variants that are associated with health conditions in ClinVar archive (ClinVar SNPs) and genome-wide association study (GWAS SNPs) between SV hotspots and the matched non-hotspot intervals (**Table S3**). Despite that there are similar amounts of coding sequences in the SV hotspots and the matched intervals, we observed a significant depletion of ClinVar SNPs (**p=0.003813, Mann-Whitney U test**) and GWAS SNPs (**p=0.002612, Mann-Whitney U test**) in hotspots as compared with the matched non-hotspot intervals. The observations that the disease-associated markers are depleted in the SV hotspots are consistent with the evolutionary scenario we proposed, that the negative selection has prevented SV hotspots at regions linked to fitness.

Majority of GWAS SNPs contribute to complex diseases in an incremental fashion (i.e., with relatively low effect sizes) and they are often under weak negative selection. The depletion of GWAS SNPs in SV hotspots suggests that the variation in these loci, in general, do not usually confer to susceptibility to complex diseases. However, it is important to make two notes here. First, given the difficulty to confidently establish the linkage disequilibrium relationship within a SV hotspot, it is plausible that some of the genome-wide association studies are underpowered to interrogate hotspots regions. Specifically, the higher repeat content of SV hotspots complicates mapping of short reads. In addition, higher levels of gene-conversion events may be expected, where linkage disequilibrium is lower than the genome-wide average, complicating imputation of individual SVs. Second, as mentioned above, some SVs in these hotspots have previously been associated with complex diseases (Sekar et al. 2016; Boettger et al. 2016; Leffler et al. 2017). As such, we argue that careful evolutionary categorizations of SV hotspots, as we are attempting here, is an important step towards understanding the complete biomedical impact of these regions.

Functional enrichment analyses can help identify clinically relevant SV hotspots. For example, one result of the GREAT analysis is that the hotspots are enriched for genomic regions linked to idiopathic bone development of distal limbs within the context of a series of *SHOX* deficiency disorders, ranging from nonspecific short stature to severe conditions such as Leri-Weill dyschondrosteosis and Langer mesomelic dysplasia (**Table S5**). When we investigated this locus closely, we observed that *SHOX* gene, deletion of which is causal to the described disorders, is located in between two individual hotspot regions (**Figure 6A**). When we overlay the breakpoints of *de novo* deletions involving *SHOX*, we found a curious clustering of breakpoints within the hotspots (**Figure 6A**).

**Figure 6.**
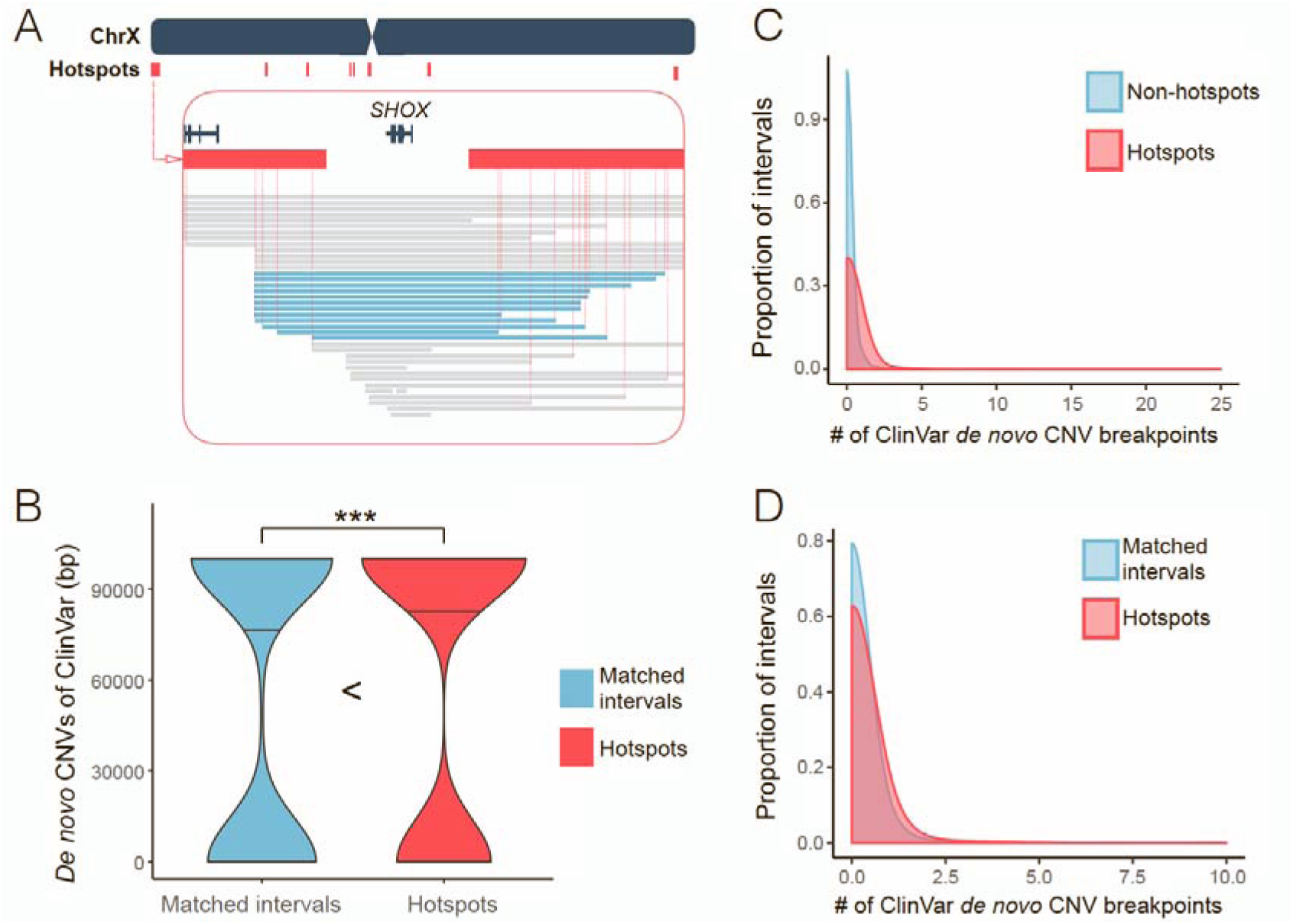
The SV hotspots and the *de novo* CNVs. (A) is a snapshot of the SV hotspots around the *SHOX* gene on chrX. There are two consecutive SV hotspots at the telomeric side of *SHOX* gene and three consecutive SV hotspots at its centromeric side (red bars). The deletions found in *SHOX* deficiency patients are represented as blue bars. There are twelve recurrent deletions spanned the hotspot regions on each sides of the *SHOX* gene. Note that the breakpoints of these deletions are estimated as the midpoints of nearby mapping markers. The exact breakpoints may be more variable. (B) shows the number of base pair(s) affected by *de novo* CNVs from the ClinVar data in SV hotspots (red) and their matched intervals (blue). The dump-belled shape of the violin plots implicate that a majority of ClinVar *de novo* CNVs are probably larger than a single interval. The *** signs indicate where there is a significant difference with a P-value lower than 0.05, calculated by Mann-Whitney U test. (C) overlays the density plots of the number of *de novo* CNV breakpoint(s) coincides with each of the hotspots (red) and non-hotspot intervals (blue). The enrichment of *de novo* CNV breakpoints in SV hotspots has a P value of 1.136×10^−5^, calculated by Mann-Whitney U test. (D) overlays the density plots of the number of *de novo* CNV breakpoint(s) coincides with each of the hotspots (red) and non-hotspot intervals with matched segmental duplication content (blue). The enrichment of *de novo* CNV breakpoints in SV hotspots has a P value of 0.008582, calculated by Mann-Whitney U test.

Previous studies found that various extremely large (generally at megabase range), *de novo*, copy number alterations (*de novo* CNVs) with similar but distinct breakpoints can lead to such disorders (Binder and Rappold 2005; Benito-Sanz et al. 2006; Chen et al. 2009). The usually devastating effects of such *de novo* CNVs prevent them to be passed down to offsprings, and therefore often discussed within the context of their recurrent mutation.

The notion that SV formation hotspots form points of plasticity in the genome that leads to the formation of large, *de novo* CNVs is an intriguing one and it also fits well with the findings of several previous studies that the segmental duplication-rich regions overlap more than expected by chance with such *de novo* CNVs (Sanders et al. 2011; Sharp et al. 2006; Xu et al. 2008; Lupski 2007). As such, we hypothesized that genome plasticity evident in SV hotspot regions may be linked to large chromosomal *de novo* events that lead to drastic developmental and cognitive disorders. Indeed, we found the hotspot intervals overlap significantly more with *de novo* CNVs from ClinVar database as compared with intervals with matched segmental duplication, exonic content, SNP, GC, and repeat elements (**Figure 6B, p=0.03929, Fisher exact test**).

This result is particularly relevant as it makes a connection between natural variation among healthy individuals and clinically relevant large *de novo* CNVs. The implication being that the plasticity observed in the SV hotspots predispose these regions to larger copy number alterations with major disease relevance. One mechanistic explanation for this observation is that concurrent mutational events in two distinct hotspot regions may lead to a large duplication or deletion event spanning the larger interval in these two hotspots. We tested this hypothesis and found that approximately ~12% of the breakpoints of *de novo* SVs (129 out of 1055 in ClinVar database) coincide with hotspot intervals significantly more than expected by chance (**Figure 6C**, **Table S6, p=1.136×10^−5^, Mann-Whitney U test)**.

This notion has been touched on in previous studies (Varki et al. 2008). We also asked whether this coincidence is primarily driven by segmental duplication content that has already been shown independently for both hotspots and for *de novo* CNV breakpoints. To our surprise, we found that the *de novo* CNV breakpoints occur in SV hotspots more often than in the intervals with matched segmental duplication contents (**Figure 6D**, **p=0.008582, Mann-Whitney U test)**, suggesting the co-occurrence of hotspots and *de novo* CNV breakpoints is not completely dependent on segmental duplication content. Overall, our results suggest that hotspots indicate plastic regions of the genome where recurrent copy number changes can emerge more often than other parts of the genome. This fits well with the earlier observation that replication stress can lead to copy number changes in the same locations where both polymorphic, as well as *de novo* events are observed (Arlt et al. 2009).

## Discussion

Genomic structural variants have increasingly been under scrutiny both within a biomedical and evolutionary context. Here, we reaffirmed previous studies that SVs are not uniformly distributed across the genome, instead they cluster within formation hotspots. Our work has attributed various architectural and evolutionary factors to the SV hotspots. We highlighted the importance of segmental duplication, relaxed negative selection, and balancing selection as the general factors associated with SV hotspots. However, we can not oversimplify all SV hotspots as results of these factors. For example, approximately two thirds of the SV hotspots do not contain any segmental duplications, and thus cannot be explained by segmental duplication mediated NAHR. The role of other SV formation mechanisms, such as double-stranded break repair and template-switching (Hastings et al. 2009) remains fascinating areas of inquiry.

Understanding the mechanistic basis of SV hotspot formation is biomedically important. In this study, we showed that hotspot regions are enriched for large *de novo* CNVs, which are often clinically relevant. These SV formation hotspots may mark regions where the likelihood of abnormal chromosomal alterations is mechanistically increased. Similar observations have been made, especially within the context of segmental-duplication mediated NAHR (Sharp et al. 2006; Bailey and Eichler 7/2006). Our results extend this observation to other mechanisms and showed that density of benign, smaller SVs can predict the breakpoints of pathogenic, larger *de novo* SVs.

### A note on background selection and SVs

The finding that NH mechanisms explain majority of the SVs in the hotspot region raises a curious issue. NH mechanisms do not heavily depend on homology or other sequence motifs. Therefore, relaxation of negative selection, rather than the genomic architecture, may be the primary driver of the formation of SV hotspots. This interpretation is further supported by the enrichment of single nucleotide polymorphisms and polymorphic mobile element insertions in SV hotspots. To follow this thread, we can conclude that non-hotspot regions of the genome may have a lower number of SVs because of negative selection. However, if that is indeed the case, the SV formation rate may actually be faster than observed but remains hidden because it would be quelled by negative selection in most of the genome.

We can envision two other forces that may contribute to the increased SNP and mobile element polymorphisms in SV hotspots. First, it is known that some single nucleotide variations can occur during SV formation, increasing the SNP content in SV rich regions (Abyzov et al. 2015). We argue that this is a small contribution and it does not explain the increased mobile element polymorphisms in the region. Second, if indeed a considerable number of SV hotspots have been maintained under some type of balancing/diversifying selection, then the increased variation is expected as is observed and reported for HLA locus (Lenz et al. 2016). Indeed, when we compared the SV hotspots that harbor genes evolving under balancing selection with other SV hotspots, we see a dramatic increase in SNP and mobile element insertion polymorphisms (**Figure S5**). As such, at least some co-occurrence of different types of variation (SV, mobile element, and SNP) can be explained by action of balancing or diversifying adaptive forces. Regardless, our study provides a strong rationale for future studies to model the background selection in the human genome that incorporates the effects of SVs.

One of the novel insights we gained from this study is that balancing selection is an observable force that maintains a subset of SV hotspots. It has been shown that some SVs can be maintained for hundreds of thousands of years through balancing selection (Pajic et al. 2016; Gokcumen et al. 2013b; Lin et al. 2015; Hollox and Armour 2008). We have now shown that some SV hotspots are also hotspots for balancing and diversifying forces. In addition to the hemoglobin clusters that we discussed above, we should also comment on the HLA locus, which is a SV hotspot not only in humans but all primate species (Gokcumen et al. 2011). At least some variation in this locus has been evolving under diversifying selection (Prugnolle et al. 2005; Yawata et al. 2006; Parham et al. 1989; Lenz et al. 2016), where different haplotypes interact with each other to form various levels of fitness effects (Lenz et al. 2015). Moreover, the *HLA* locus in humans is enriched for Neanderthal haplotypes (Abi-Rached et al. 2011) and *HLA* variation within specific African populations is adaptively shaped by admixture events (Patin et al. 2017). As such, we argue that we can conceptualize the *HLA* locus as a reservoir of variation, SV and single nucleotide variation alike, where diversity is adaptively maintained through rapid changes in pathogenic pressures, and different fitness effects of diverse HLA haplotype combinations. Such adaptive scenarios can also apply to other SV hotspots, including those that harbor immune-related SV hotspots, including killer-cell immunoglobulin-like receptor genes (Traherne et al. 2010; Jiang et al. 2012; Pelak et al. 2011), immunoglobins (Watson et al. 2013), and defensin genes (Aldred et al. 2005).

Another related adaptive scenario within the context of SV hotspots is convergent balancing selection. There are few such examples in humans where recurrent mutations with similar phenotypic effect are maintained in different populations through balancing selection (Sweeney et al. 2017). In fact, some of the variation in the *HLA* locus has been suggested to evolve in such a manner (Titus-Trachtenberg et al. 1994; Erlich and Gyllensten 1991). Our example of the globin loci, where loss of function mutation due to deletions can lead to resistance to malaria with the cost of causing blood disorders (**Figure 5**) also fit in this general category. Another recent example is the *GYPB* locus, which is an SV hotspot. Leffler et al. showed geography-specific selection for a given SV in this region, as this SV provides resistance to malaria a similar situation with what is observed for globin loci (Leffler et al. 2017).

In sum, our results identify a subset of SV formation hotspots, which are enriched for genes that are involved in environmental interaction, and implied in multiple human diseases. It seems that diverse forms of balancing selection (Key et al. 2014) maintain the variation in these hotspots. The characterization of SV hotspots also give hints about the global adaptive and mechanistic forces that shape the human genome. Our work follows previous work that investigates the genomic and biomedical impact of SVs using evolutionary framework. For example, Makino et al. highlighted the importance of cross-species SV deserts in identifying disease-related SVs in humans (Makino et al. 2013). Rice and McLysaght, as well as Makino et al. used conservation, both at the sequence and copy number levels, as a proxy to show that SV pathogenicity can be predicted by dosage sensitivity (Rice and McLysaght 2017). In sum, by constructing an updated map of SV hotspots, and outlining their broad evolutionary implications, our study opens new avenues for research to better understand genome evolution and biomedical implications of complex variations.

## Methods

### Data Sources

We obtained the data used in this project from the following sources: The sequencing gaps were from Genome Reference Consortium (International Human Genome Sequencing Consortium 2004). The segmental duplications were from Bailey at el. (Bailey et al. 2002). The fixed mobile elements were from RepeatMasker (Smit et al. 2016). The SNPs information were from dbSNP (Sherry et al. 2001). The coding and genic sequences were from NCBI RefSeq Project (Pruitt et al. 2014). The GWAS SNPs information was from GWAS catalogue (Hindorff et al. 2009). The ClinVar variants were from NCBI ClinVar archive (Landrum et al. 2016). The DNase I hypersensitive sites were from ENCODE (ENCODE Project Consortium 2012). The above data were available through UCSC Table Browser (Karolchik 2004).

We parsed the data from 1000 Genomes Project Phase 3 release (1000 Genomes Project Consortium et al. 2015) for the coordinates of genomic structural variants and polymorphic mobile elements. The formation mechanisms of a subset of SVs were available from Abyzov et al. (Abyzov et al. 2015). The B-statistics value were available from McVicker et al. (McVicker et al. 2009). The list of genes under balancing selection were available from DeGiorgio et al. (DeGiorgio et al. 2014). The methodologies used to minimize the false positive calls for balancing selection, such as avoiding sites overlapping SVs as well as sites that show excess heterozygosity (indicating mapping error), make this list particularly conservative in SV-rich regions. The deletions discovered from *SHOX* deficiency patients were available from Benito-Sanz et al. (Benito-Sanz et al. 2006). We estimated the breakpoints of these deletions as the midpoints of the mapping markers.

### Hotspot determination

We divided the autosomes and X chromosome of human genome (hg19) into 100kb intervals. To ensure that all intervals in our dataset have the same length of annotated sequences, the remainder sequences at the distal end of chromosomes which do not make up to a full-length interval (23) are discarded from our dataset. Similarly, the intervals overlapping with sequencing gaps (2,250) are discarded from our dataset. Finally, we created a dataset with 28,103 intervals.

We mapped the 42,758 SVs (deletions, duplications, multiallelic copy number variants, inversions, and insertions) from 1,000 Genomes Project Phase 3 data release to the 28,103 intervals from our dataset. We assumed that the number of SVs hitting each interval would follow a Poisson probability distribution if SV distributed randomly across the genome, and built the expected Poisson distribution using the average number of SVs in each interval. The expected Poisson distribution was used to decide the criteria for SV hotspots. The intervals with a SV number equaling to or higher than the 99 percentile of the estimated Poisson probability distribution are determined as SV hotspots.

### Genomic content measurement

We used bedtools (Quinlan and Hall 2010) to quantify the overlapping of genomic features to each of the intervals in our dataset. We quantified the coverage of segmental duplications, coding sequences, genic sequences, fixed mobile elements, SNPs, GWAS SNPs, *de novo* SNPs, *de novo* CNVs, and DNase I hypersensitive sites in each interval for further comparison. We quantified the number of genes under balancing selection, SVs with specific types, and polymorphic mobile elements that have any overlap with each interval. We quantified the B-statistics of an interval by summing up the B-statistics value of all sites of the interval and dividing it by the total length of an interval.

### Matching intervals

For each SV hotspot, we used a customized python code (available through github, https://github.com/ontalin/matching-intervals) to screen out all the non-hotspot intervals that have the required genomic content similar to the hotspot (10% differences is tolerated) and randomly pick one from the subset as its matched non-hotspot interval.

### Ontology enrichment analysis

We used the “two unranked lists of genes” mode of GOrilla to perform GO overrepresentation test of genes (Eden et al. 2009). Specifically, we built a list of hotspot genes with RefSeq genes that have at least one base pair of their coding sequences overlapping with SV hotspot, and a list of background genes with RefSeq genes that have at least one base pair overlapping with any interval of our dataset. The target list was compared against the background list for overrepresented GO terms. The GO terms with their FDR q-value smaller 0.05 are considered significantly overrepresented among the SV hotspot genes.

We used GREAT (McLean et al. 2010) to perform ontology overrepresentation test of cis regulatory sequences. We used the bed file of SV hotspots as the target regions and the bed file of all intervals in our dataset as the background regions.The ontology terms with Hyper FDR Q-Value smaller than 0.05 are considered overrepresented among the SV hotspots.

## Acknowledgements

We like to thank our friends in the Gokcumen Lab: Marie Saitou, Ozgur Taskent, Petar Pajic, and Izzy Starr for careful reading of this manuscript and multiple discussions to better present the data. We also are grateful to our colleagues Amanda Larracuente, Trevor Krabbenhoft, and Derek Taylor for their insightful criticism, which improved our manuscript substantially. Last, We want to acknowledge The National Science Foundation under (Grant No. 1714867) that allowed us to expand on our observations in the epidermal differentiation complex to genome-wide studies.

